# Aged mouse ovarian immune milieu shows a shift towards adaptive immunity and attenuated cell function

**DOI:** 10.1101/2021.08.12.456051

**Authors:** Tal Ben Yaakov, Tanya Wasserman, Yonatan Savir

## Abstract

The immune system plays a major role in maintaining many physiological processes in the reproductive system. However, a complete characterization of the immune milieu in the ovary, and particularly how it is affected by maternal aging, is still lacking. In this work, we utilize single-cell RNA sequencing and flow cytometry to construct a complete description of the murine ovarian immune system and its changes along with pre-estropause aging. We show that the ovarian immune cells composition undergoes an extensive shift with age towards adaptive immunity. We analyze the effect of aging on gene expression and chemokine and cytokine networks and show an overall decreased expression of inflammatory mediators together with an increased senescent cells recognition. Our results reveal the changes in the aging ovarian immune system of the fertile female as it copes with the inflammatory stimulations during repeated cycles and the increasing need for clearance of accumulating atretic follicles.

## Introduction

One of the effects of aging in mammalians is a decline in fertility and hence a diminished capability to give birth to offspring^1^. In women from their early 30’s, there is a sharp decrease in fertility, accompanied by an exponentially increase in the odds of miscarriages and birth defects, alongside a drastically lower success rate of in-vitro fertilization (IVF) procedure^2–4^. Another well-documented effect of age is on the ability of the immune system to overcome illnesses and eliminate different pathogens^5–8^.

The presence of immune cells such as macrophages (M*ϕ*s), dendritic cells (DCs), granulocytes, T and B lymphocytes was identified through the entire female reproductive tract^9,10^, and in the ovaries in particular^11–14^. These cells participate in many fertility-related processes in the ovaries – from follicle development up to ovulation and corpus luteum formation and regression^13,15–18^. Ovulation, for example, is considered an inflammatory process that includes edema, vasodilation, pain, and heat^19,20^. Changes in the immune milieu, such as depletion of M*ϕ*s and DCs have been shown to result in a decreased number of ovulated oocytes, depletion of endothelial cells, increased follicular atresia, and lead to a delayed progression of the estrus cycle^16,17,21^.

Characterizing the complete immune milieu in the ovary, and in particular how it is affected by maternal aging, is challenging mainly due to the small fraction of the immune cells compared to other cell types in the ovary^22,23^. For that reason, previous work using whole-ovary single-cell measurements did manage to portray the ovary’s main cell types (oocytes, granulosa, theca, immune, etc.) yet didn’t have the resolution to resolve the entire immune milieu^22–24^. Others have a priori focused on a limited set of cell types^16,17^, while additional studies have used bulk RNA sequencing experiments^25^, hence did not capture the entire immune members in the ovaries. Attempts to characterize the effects of maternal age on the immune system in the ovary were therefore limited to a small subset of cells^23,26^, addressing mainly changes in the macrophages fraction that decreases with age, and accumulation of inflammatory mediators such as cytokines and reactive oxygen species within the tissue^23,26^.

In this work, we provide, for the first time to our knowledge, a complete detailed characterization of the murine ovarian immune system composition at the single-cell level. We show the presence of various immune cell populations, such as M*ϕ*s, DC’s, neutrophils (NTs), NK cells, NKT cells, innate lymphoid cells (ILCs), B cells, and several T cell types – including an ovary specific CD3^+^ CD4^−^ CD8^−^ double-negative T (DNT) cells. Moreover, we show an extensive tissue-specific effect of maternal age on the ovarian immune milieu, resulting in a shift towards adaptive immunity, mainly by a significant increase in the DNT population. In addition, we analyzed the changes in gene expression of the cells and discovered a global attenuation in their general function and responsiveness. We also found a decrease in the expression of inflammatory mediators such as cytokines and chemokines. Moreover, we identified some evidence for an increase in senescent cells recognition activity. Our results serve as an opening for a much more comprehensive understanding of the interaction between maternal aging and the immune system in fertile female mammals.

## Results

### The ovarian immune milieu consists of various cell types and alters at old age

To characterize the ovarian immune milieu, we have isolated immune cells (CD45^+^ cells) from the ovaries of young (11-15 weeks), adult (20-37 weeks), and old (40-47 weeks) virgin mice, and utilized flow cytometry and single-cell RNA sequencing (scRNA-seq) to characterize the cell types and how they are affected by maternal age (Fig. 1a). First, we performed scRNA-seq on isolated immune cells from the ovaries of 13-weeks old mice (n=2, 3,307 cells). To cluster the cells and identify their type, we used a combination of both literature-based annotation and automatic annotation methods (Seurat R package and SingleR algorithm, “Methods” and Supplementary Fig. 1). The combination of these methods allowed us to identify within the ovaries the following cell types: Mφs, DCs, NTs, B cells, NK cells, NKT cells, ILC1, ILC2, ILC3, and several clusters of T lymphocytes: CD8^+^, CD4^+^, and CD4^−^ CD8^−^ double-negative T cells (DNT cells) (Fig. 1b,c). The majority of the cells were innate immune cells, mostly ILC1, Mφs, NTs, and NK cells. To validate the frequency of group 1 of innate lymphoid cells (G1-ILC) in the ovaries, we conducted a flow cytometry experiment measuring the fractions of CD45^+^ NK1.1^+^ CD3^−^ cells (i.e. NK and ILC1) in the mouse ovaries and spleen (Supplementary Fig. 2). The average fraction of G1-ILC in the spleen was 11.81% and in the ovary was 42.71%. These results are consistent with the high G1-ILC fraction resulting from the ovary scRNA-seq analysis and previous results, which demonstrated that G1-ILC proportion in mice spleen is relatively low^27^. Among other cell types that were found, DNT cells are unique, somewhat less well-defined cell population. To confirm the presence of CD4^−^ CD8^−^ cells in the ovaries, we conducted a flow cytometry experiment comparing the fractions of CD4^+^, CD8^+^, and CD4^−^ CD8^−^ cells in the mouse ovaries, spleen, and peritoneum (Supplementary Fig. 3). These measurements validate the scRNA-seq results and show that although present in other tissues at small fractions, CD3^+^ TCRβ^+^ CD4^−^ CD8^−^ cells are tissue-specific cells to the ovaries.

**Figure 1.**
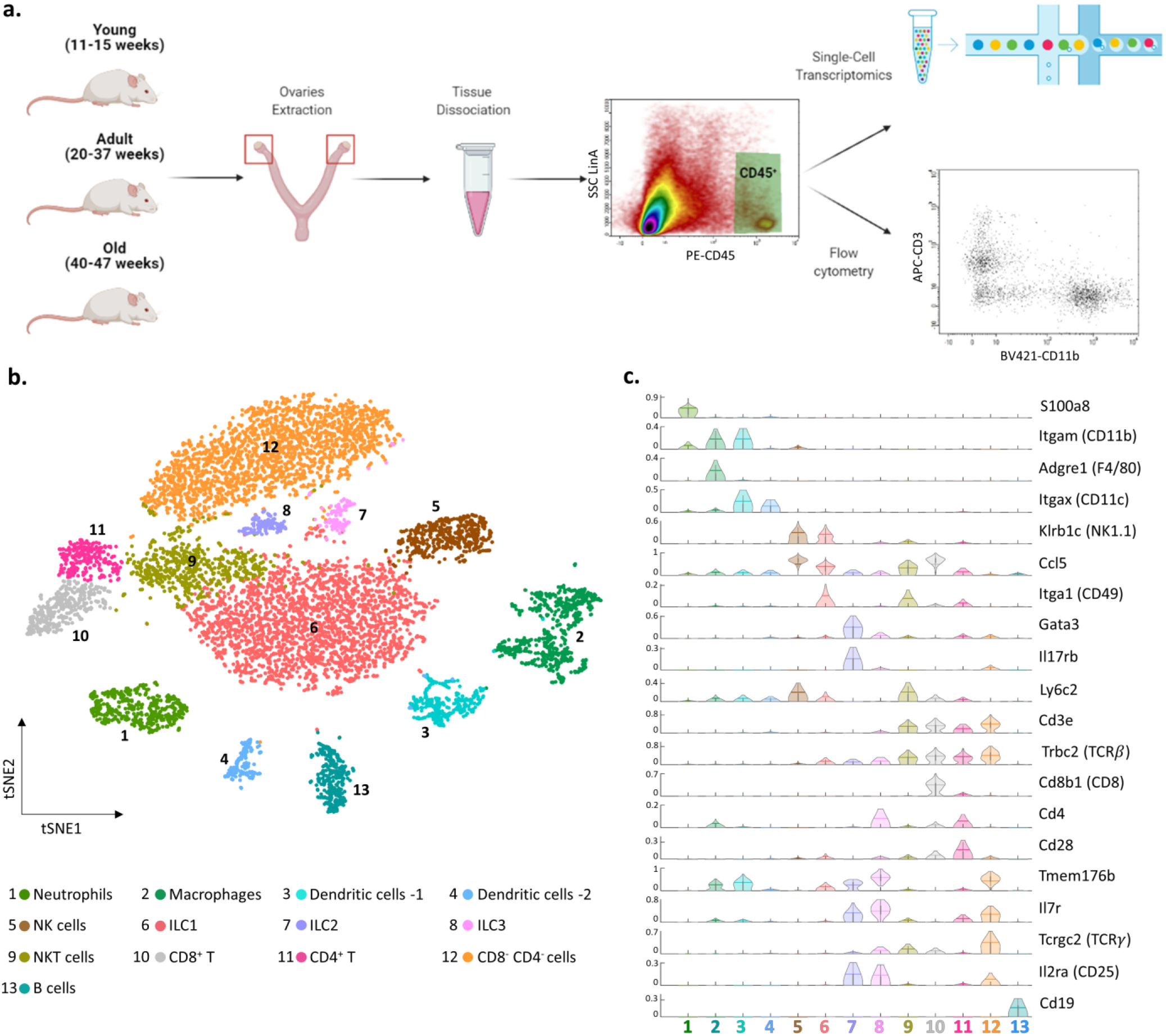
The ovarian immune milieu is consisted of various cell types. (a) Schematic illustration of the experimental pipeline (created with BioRender.com). Ovaries of female mice at different ages were extracted. Then, cells were gated for CD45 expression and further analyzed using single-cell RNA sequencing or flow cytometry. (b) tSNE plot of joint data from both samples (young and old), divided into clusters. (c) Violin plot of normalized expression for cluster-specific markers. Each row represents the normalized expression of a single marker across all immune clusters. Normalized expression values are between 0 to 1.

**Figure 2.**
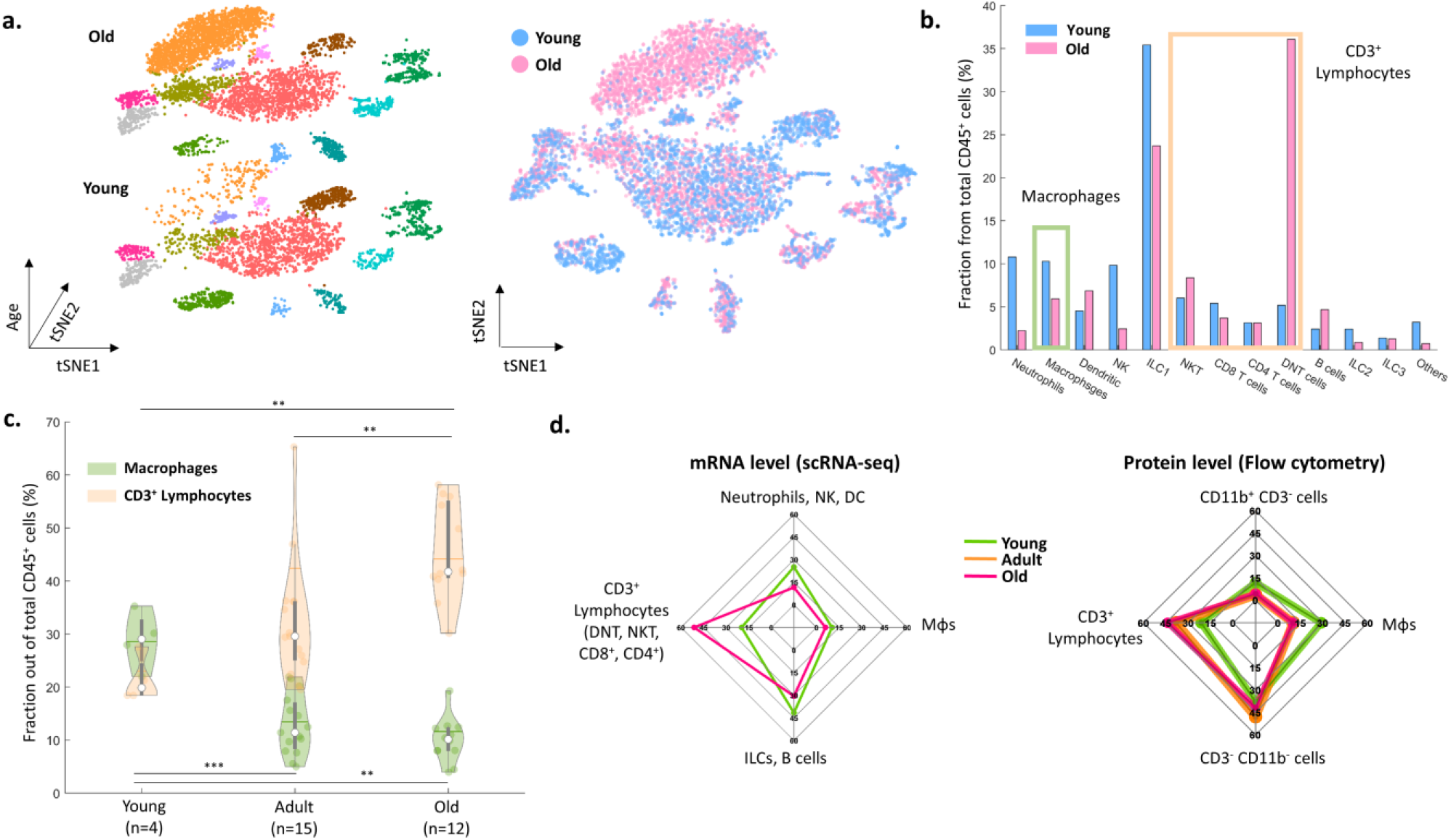
The effect of maternal age on the ovarian immune milieu. (a) 3-D tSNE plot (left) and an overlay (right) of all ovarian CD45^+^ cells found in scRNA-seq, divided by age group. (b) The effect of maternal age on the fractions of each cell type. The green and yellow rectangles mark the macrophages and CD3^+^ populations, respectively. (c) Violin plot of the changes in fraction distributions of macrophages and CD3^+^ lymphocytes as a function of age as measured by flow cytometry (Kolmogorov-Smirnov test, ** p value<10^−2^, *** p value<10^−3^). (d) Comparison between transcriptome and protein level of immune populations within the ovaries at different maternal ages. Each spider plot shows the distribution of different immune cell types measured using scRNA-seq (left panel) and flow cytometry (right).

**Figure 3:**
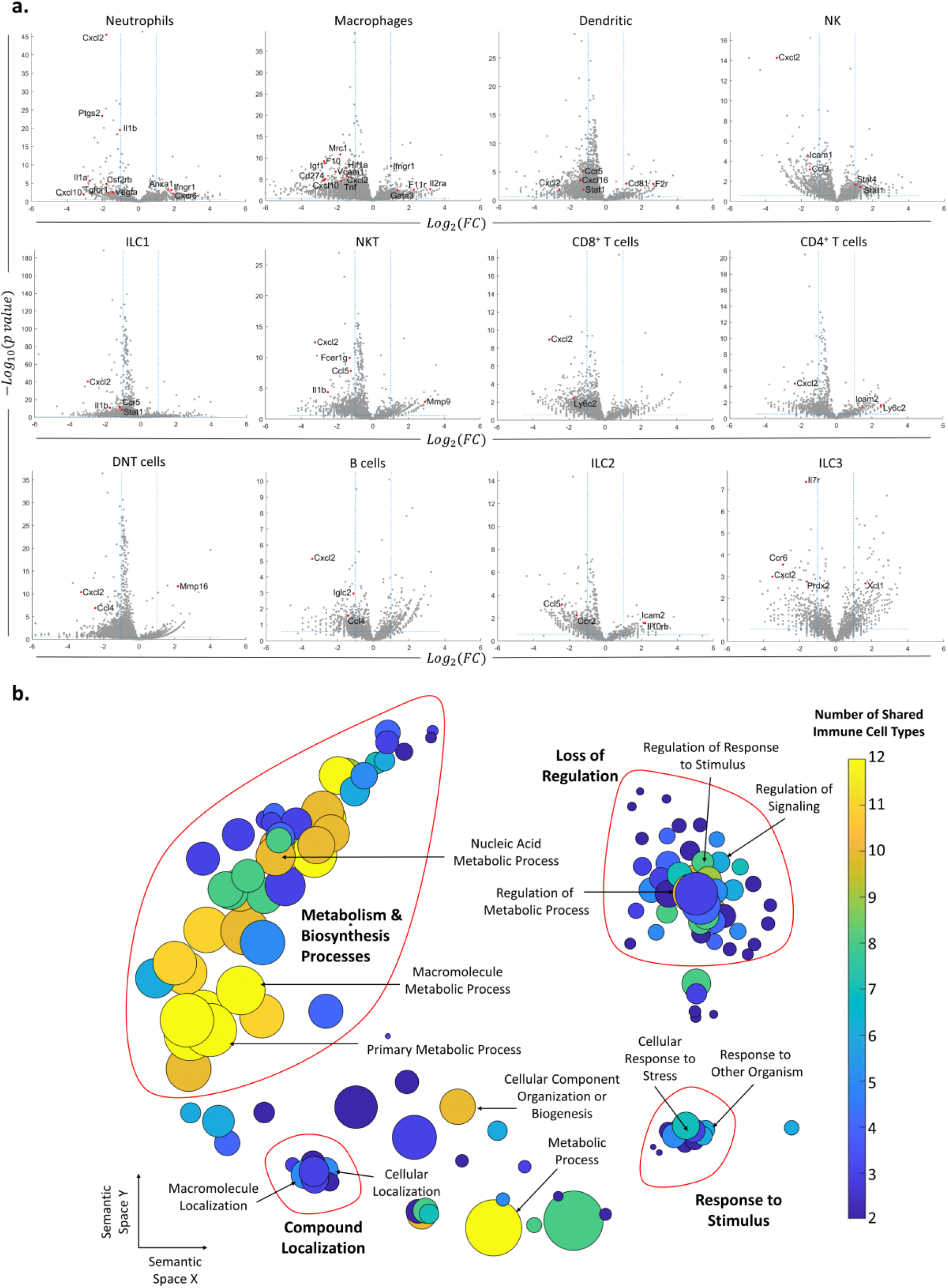
Changes in gene expression along with maternal age. (a) Volcano plots of the scRNA-seq analysis for the different immune cell types. The horizontal dashed line mark p-value=0.05, and the vertical dashed lines mark 2-fold upregulation and downregulation. Genes in the top left section in each graph are significantly downregulated at old age, while genes in the top right section are upregulated at old age. (b) Each circle represents a downregulated biological process in the old mice, which appeared in at least two types of immune cells. The axes represent semantic similarities distance as calculated by REVIGO (“Methods”). The color of each term represents the number of immune cell types in which the process is downregulated with age. The size of each term represents the hierarchy of the biological process; the bigger the circle, the higher the hierarchy of the process is.

Next, we examined the changes in the ovarian immune milieu at older ages. Using cells isolated from old, near estropause (43 weeks; the rodent equivalent of the human menopause), mouse (5,468 cells) we characterized the old ovarian immune system (using the same annotation methods) and compared it to its younger counterpart. The results demonstrated a shift at older age towards a lymphocytes-rich environment that was accompanied by decreased fractions of several immune populations such as ILC1 cells, Mφs, NTs, and NK cells (Fig. 2b). To both validate the scRNA-seq results and to check whether this effect is cycle-stage dependent, we conducted several flow cytometry experiments. To measure the effect of maternal age on the fractions of T-lymphocytes (CD3^+^) and Mφs (CD11b^+^ F4/80^+^) from total ovarian CD45^+^ cells, mice were divided into three groups of age: young (11-15 weeks, n=4), adult (20-37 weeks, n=15) and old (40-47 weeks, n=12). Results show a significant increase along with age of CD3^+^ cells’ fraction (young vs. adult and adult vs. old, Kolmogorov-Smirnov test, p<10^−2^), while the fraction of Mφs was significantly decreased (young vs. adult and young vs. old, Kolmogorov-Smirnov test, p<10^−3^ and p<10^−2^ respectively) at older ages (Fig. 2c,d). Moreover, these age-dependent results are cycle-stage independent (Supplementary Fig. 4). Taking the results both from the scRNA-seq the flow cytometry ones (Fig. 2d), there is a consistent shift towards adaptive immunity (an increase in the CD3^+^ lymphocytes fraction), while most innate immune cells’ fraction (Mφs, NK, ILC1, and NTs) decreases.

**Figure 4.**
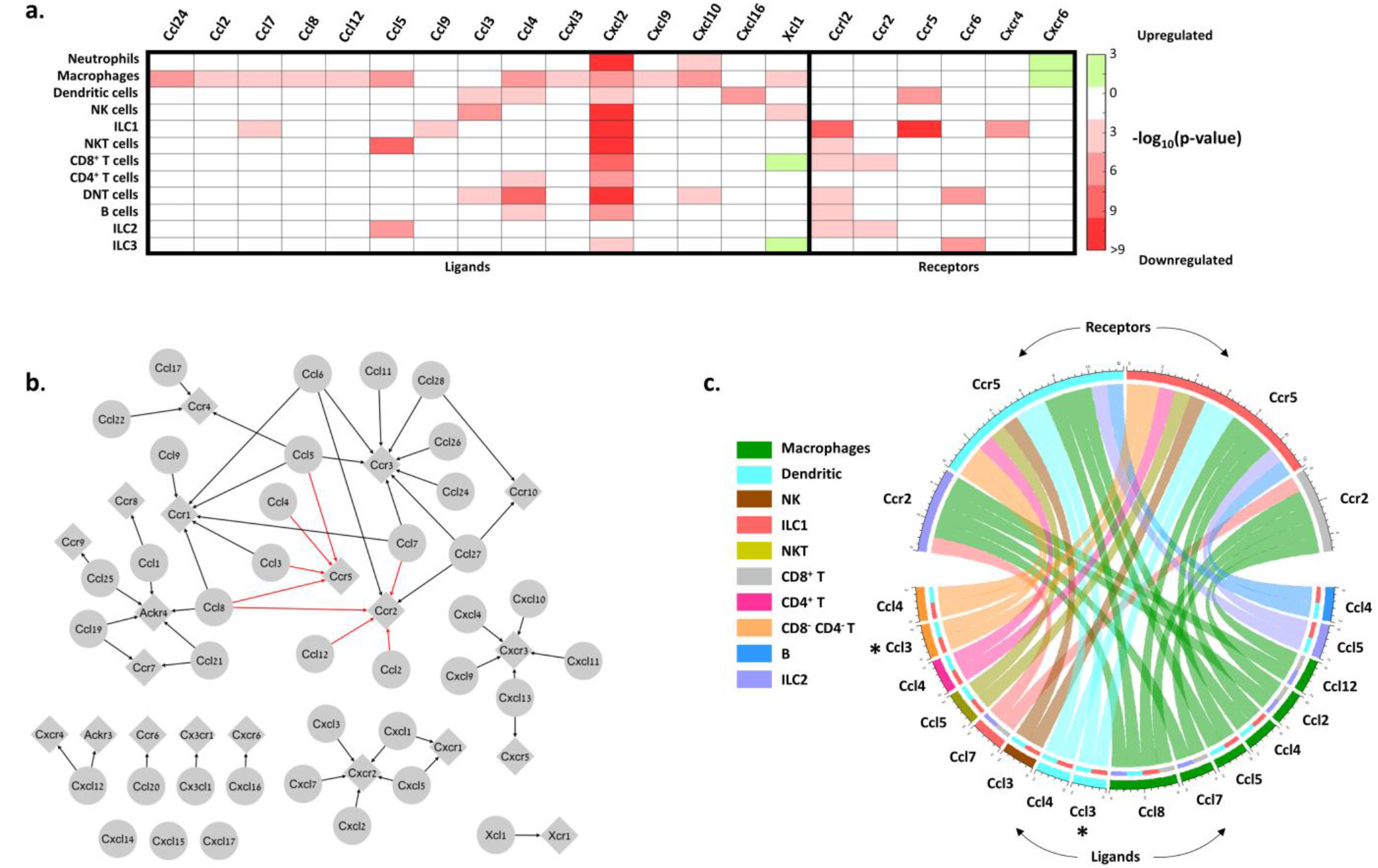
The effect of maternal age on the chemokines network. (a) A heat map of significantly (p<0.05) 2-fold decrease (red) or increase (green) in the expression of chemokines and chemokine receptors genes in different immune cell types. (b) The downregulation of the chemokines network due to age. Significantly (p<0.05) downregulated edges are highlighted in red. (c) Chord diagram of downregulated edges in the chemokines interaction network. The colors in the outer circle denote the immune cell type expressing the chemokine or receptor, while the chord color is the same as the immune cell type expressing the ligand node of each edge. For example, the asterisks denote the downregulation of the edge that connects *Ccl3* in DNT or DCs and *Ccr5* in ILC1 and DCs.

### The effect of maternal aging on the ovarian immune cells’ transcriptome

After identifying the ovarian immune milieu and the changes it undergoes at older age, we characterized the changes in gene expression within each immune cluster. Figure 3a depicts the differentially expressed genes patterns across all clusters. Several clusters, such as DNT cells, ILC1, NKT cells, and CD4^+^ T cells exhibited an extremely skewed pattern, in which most of their differentially expressed genes (DEGs) were downregulated, compared to upregulated DEGs. This may imply that these cell types are more susceptible to maternal aging. After defining DEGs for each cell type, we explored changes in biological processes via GO enrichment analysis (Fig. 3b, “Methods”). Most cell types showed an enriched set of processes that were downregulated with maternal age. Using the REVIGO platform (“Methods”), we eliminated redundant GO terms and counted the appearances of each GO term across all cell types (Supplementary Table 1). Terms that were found in more than one cell type were classified as “global”, and cell type-unique terms as “specific”. For further analysis, we took all the global terms and used the REVIGO platform to cluster them according to their semantics distance (Fig. 3b). The results show a global decrease in several clusters of processes. The clusters with the processes that were mostly shared among cell types included decreased biosynthesis and metabolism-related processes. Another distinct cluster includes a general loss of regulation over various processes. The third well-defined cluster includes decreases in the cellular response to different stimuli.

Among the cell-type-specific downregulated processes, Mφs exhibit attenuation in immune and inflammatory responses, along with decreased tissue remodeling and wound healing processes. DCs showed a decrease in cell activation and regulation of immune response. Several immune cell types showed a limited set of enriched upregulated processes, in which the most prominent ones were exhibited by Mφs and B cells and included T cell activation and differentiation processes (Supplementary Table 2).

### Aging affects the cytokines and chemokines connectome of the ovarian immune cells

To get a better notion of the effect of aging on the different immune cell types, we estimated the effect of aging on the chemokine and cytokines interactions between the immune cells (Fig. 4 and Fig. 5). For both chemokines and cytokines, we used the KEGG database to extract the network of ligands and receptors and their interactions (“Methods”). Within each network, we focused on significantly changed connections, which we defined as edges that both of their nodes (i.e. both the ligand and receptor) have a significant (p-value < 0.05) 2-fold or higher decrease or increase in their expression at old age. We found that almost all significant nodes and edges in both networks were downregulated (Fig. 4a,b and Fig. 5a,b).

**Figure 5.**
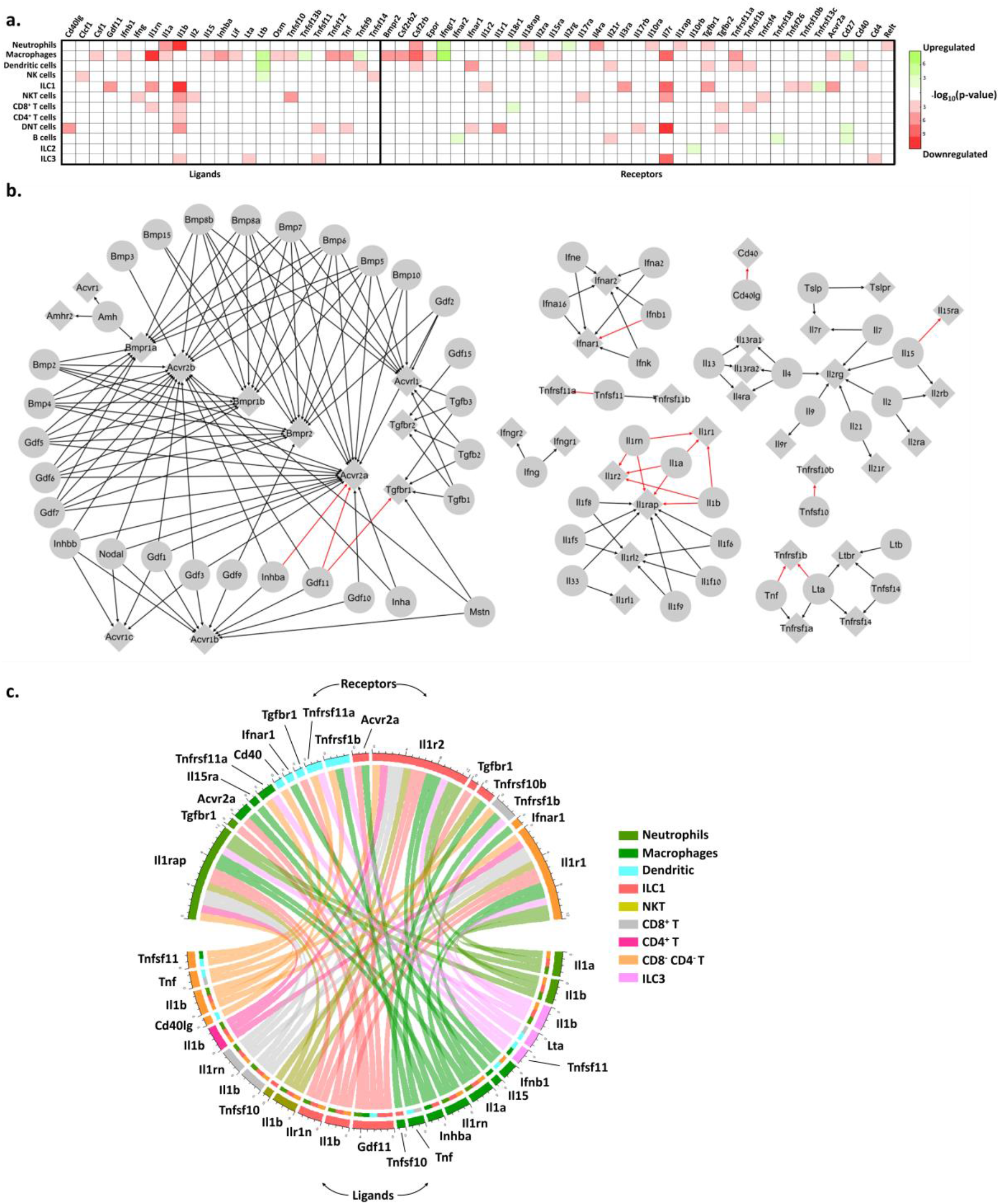
The effect of maternal age on the cytokines network. (a) A heat map of significantly (p<0.05) 2-fold decrease (red) or increase (green) in the expression of cytokines and cytokine receptors genes in different immune cell types. (b) The downregulation of the cytokine network due to age. Significantly (p<0.05) downregulated edges are highlighted in red. Only the connected regions of the network that have downregulated edges, in addition to the IFNγ subgraphs, are shown. (c) Chord diagram of downregulated edges in the cytokine interaction network. The colors in the outer circle denote the immune cell type expressing the cytokine or the receptor, while the chord color is the same as the immune cell type expressing the ligand node of each edge.

The most prominent affected edges in the chemokines network involved *Ccr5* expressed by DCs and ILC1 cells, *Ccr2* expressed by CD8^+^ T cells and ILC2 cells, and their ligands *Ccl2, Ccl3, Ccl4, Ccl5, Ccl7, Ccl8*, and *Ccl12* (Fig. 4b). Both of these receptors have been shown to take part in chemoattraction of immune cells in the context of various inflammatory processes^28–30^. Moreover, *Cxcl2*, an inflammatory chemokine that mediates neutrophils trafficking^31,32^, was significantly decreased in almost all immune cell types (Fig. 4a). CCRL2 is an atypical chemokine receptor that was found to be upregulated in activated immune cells after induction of inflammatory signals^33^. *Ccrl2* was downregulated in several cell types such as ILC1 and NKT, although it is mainly expressed by myeloid cells such as NTs, DCs, and Mφs^33^.

The changes in the cytokines network were found mainly in the IL-1 superfamily (*Il1r1, Il1r2*, and *Il1rap* along with *Il1a* and *Il1b –* but also the *Il1rap* antagonist, *Il1rn*). Moreover, TNF-receptor *Tnfsfr1b* and its ligands *Tnf* (TNF*α*) and *Lta* (TNF*β*) also showed a significant decrease at old age (Fig. 5). As IL-1 and TNF superfamily members are considered inflammatory, along with the evident decrease in various inflammatory chemokines and their receptors, our results suggest that beyond overall inhibition in the ovarian immune’s function, aging also shifts its phenotype towards a less inflammatory state.

Another evident downregulated edges in the cytokine network were in the TGF*β* superfamily (*Gdf11* and *Inhba* along with their receptors *Acvr2a* and *Tgfbr1*). Activin A, a dimer composed of two Inhibin-*βA* subunits (the translation product of *Inhba*), is produced among others by the gonads and promotes LH secretion from the pituitary. It plays an important role in expanding the primordial follicle pool and contributes to the early stages of follicular growth by increasing FSH receptor expression on granulosa cells^34^. In addition, Activin A was found to activate resting macrophages – yet there are contradictory findings as to rather its effect is pro or anti-inflammatory^35^. Decreased expression of both *Inhba* and *Acvr2a* by ovarian macrophages at an older age might suggest a specific role of macrophages in supporting follicular growth (via Activin secretion) during the estrous cycle, which decays as age progresses. Moreover, these results may present a mechanism in which macrophages are participating in inducing an inflammatory environment as part of the ovulation process as a response to Activin.

### Aging may affect the clearance of senescent cells by immune cells in the ovaries

Inducing cell senescence, which is an irreversible state of growth arrest, is a mechanism the body uses to handle cell stress which can accumulate during aging^36^ and may result in chronic diseases and tissue dysfunction^37,38^. One of the main molecular features of senescent cells is the senescence-associated secretory phenotype (SASP), in which the senescent cells create an inflammatory environment by secreting inflammatory cytokines, chemokines, growth factors, extracellular remodeling factors, and more^39,40^. Immune cells respond to these factors, detect specific markers expression or their absence on the senescent cells’ membrane and clear them either by phagocytosis (by Mφs, for example) or by killing (by NTs or NK cells for example)^39^. Ovarian senescence was already studied in the past^41^, however, the specific mechanism of senescent cells clearance within the ovaries is still unclear.

We found that the fraction of cells that express *Ccr2, Csf2ra*, and *Csf1r*, which are all receptors for known SASP proteins^39,42^, was significantly higher in old Mφs (Supplementary Fig. 5). In addition, the fraction of old Mφs that express cell surface markers that were previously reported to take part in the recognition of senescent cells, such as membrane IgM’s (*Ighm*) and C-type lectin receptors (*Clec4a2-3*)^43^ was also elevated. Moreover, old NTs and Mφs showed upregulated expression of *Ifngr1* (Fig. 5a), a part of the IFNγ receptor, while the cytokine itself is overexpressed by senescent cells^44,45^. In addition, both NTs and Mφs, as well as NK cells showed a higher expression fraction of this receptor. NTs cells also exhibited elevated fractions of *Ccr1*, a receptor for several SASP chemokines^46^, while old NKT cells had higher levels of *Cd74*, a receptor for MIF, another SASP member^46,47^. CXCR6 is another novel mediator of senescence control which was recently discovered as part of senescence surveillance in the liver by CD4^+^ and NKT cells^48^. *Cxcr6* expression was also increased in old NTs and Mφs.

## Discussion

In this work, we characterized for the first time as far as we know, the complete mouse ovarian immune milieu and its changes with maternal age. We have identified by both single-cell RNA sequencing and flow cytometry various immune cell types, such as macrophages, neutrophils, dendritic cells, NK cells, ILCs, NKT cells, B cells, and several T cell subtypes. We investigated the changes in the composition of the immune compartment along with age, up to near-estropause state, and discovered major changes in the fractions of different immune cell types. Most innate immunity cells, such as NTs, M*ϕ*s, ILCs, and NK cells exhibited a decrease in their fractions from the total immune population. Moreover, there was a substantial increase at an older age in the presence of predominantly DNTs (CD3^+^ TCRβ^+^ CD8^−^ CD4^−^ T cells).

Our results suggest an elevation in CD3^+^ TCR*β*^+^ lymphocytes along with age, which could not be explained solely by the elevation of CD4^+^ or CD8^+^ T cells. The results indicate a dramatic change in the fraction of DNT cells (from ~5% to ~35% at old age). Other identified DNT cells, in the blood or lymph nodes, for example, consist 1-5% out of all the lymphocytes^49,50^, which is significantly less than the fraction we found in the ovaries, regardless of age. These results are consistent with a previous study that showed an increase in the TCR*β*^+^ lymphocytes that is not due to CD4^+^ or CD8^+^ in mice before estropause^23^.

Double-negative T cells were found to be the most common lymphocyte across the female mice uterus and cervix^51^. These cells were shown to have a regulatory function, showed no proliferation, inhibited the proliferation of splenic T cells when co-cultured together *in vitro*, and their origin was suggested to be extrathymic. The DNT population we observed in the ovary, showed almost no expression of classical thymocytes development markers at the double-negative stages such as Notch1, Socs3, Dtx1, Hes1, and more^52^. Other studies showed that DNT cells in peripheral blood or lymphoid organs have a suppressive role and assist in preventing allograft rejection and autoimmune responses^50,53^. Some studies suggest that DNT cells are the result of down-regulation of CD4 and CD8 due to chronic stimulation^54,55^. The ovaries present ongoing cycles of inflammation processes in each ovulation, which may be resulted in chronic stimulation of ovarian T cells and may be consistent with an increase of CD4^+^ T-call after estropause^23^.

ILC1 cells are the largest population in the young mouse ovaries, and the second largest in the old ovaries, after the DNT population. These cells, along with NK cells, constitute the group 1 innate lymphoid cells. As opposed to NK cells, ILC1s have weak-to-no cytotoxic capabilities, and are more similar to Th1 cells in their function, i.e. inducing type 1 immune response against intracellular pathogens and inflammatory responses^56^. In addition, ILC1 cells take part in the process of tissue remodeling^57^. Interestingly, ILC1 distribution was also found to be elevated in the epididymal adipose of male mice’s testis^27^. As a dynamic, constantly changing environment, the ovaries, which involves cycles of developing and regressing structures, may use ILC1 cells as orchestrators of other cell types participating in these processes.

Taking together the observed changes in global processes, and expression of immune mediators such as chemokines and cytokines, a state of general attenuation and unresponsiveness is emerging. GO enrichment analysis of joint processes for several cell types present a decrease in general activity, in which immune cells exhibit lower levels of metabolism-related processes accompanied by decreased responsiveness to different stimuli and loss of regulation over various processes. Analysis of the chemokines network also revealed general attenuation and decreased expression of both chemokines and their receptors by different cell types, mostly of *Cxcl2* which is responsible for neutrophils recruitment, *Ccr5* signaling in ILC1 and DCs, and *Ccr2* signaling in CD8^+^ T cells and ILC2. Moreover, several inflammatory chemoattractants, such as *Ccl2, Ccl3, Ccl4*, and *Ccl5*, as well as pronounced inflammatory cytokine transcripts such as TNFα, IFNγ, IL1-α, and IL-1β were all decreased in different immune cell types. Moreover, decreased expression of several members of the TGF*β* superfamily by macrophages at old age, such as *Inhba* and *Acvr2a*, might be connected to impaired or decreased follicles growth. Our results suggest a decrease in the inflammatory characteristics of ovarian immune cells which may contribute to ovarian dysfunction, as ovulation cycles are considered as repeated inflammatory processes.

We have found an increase in genes involved in senescent cell recognition by several immune cell types, mostly M*ϕ*s but also NTs, NK, and NKT cells. Generally, as age progresses, senescent cells are accumulating within tissues^41,58,59^. In the ovaries, aging and senescence are accompanied by an increasing number of atretic follicles and cycle irregularities that end in estropause. Our results suggest that clearance of senescent ovarian cells by immune cells increases at old age to match the ovary needs.

There is a plethora of studies that found an age-related increase in the inflammatory environment in the body, a state that develops over time and terms “Inflammaging”^60–62^. Our results show that the most evident change in the ovaries of aging mice before estropause is a shift from innate to adaptive immunity, which is associated with a dramatic increase of CD3^+^ lymphocytes, and a decrease in macrophages, NTs, and ILC1 fractions. Besides the changes in the composition of the immune milieu, most cell types exhibit a decrease in inflammatory hallmarks. Our results are consistent with a previous work that shows a similar trend in mice before estropause, with elevated levels of inflammatory mediators only in mice older than one year^23^. Our results demonstrate how the fertile female’s ovarian immune system ages while coping with the two main challenges of the aging ovary before menopause: the inflammatory stimulations due to repeated cycles and the increasing need for clearance of accumulating atretic follicles.

## Methods

### Mice

All experiments involving mice conform to the relevant regulatory standards (Technion IACUC and national animal welfare laws, guidelines, and policies). Hsd:ICR female mice were purchased from Envigo RMS (Israel) and were housed in a 12h light / 12h dark cycle. Assessment of estrous cycle stage for the mice begun 3 days prior each experiment via vaginal smears^63^. In short, mice’s vaginal canals were washed using 20μL saline (PBS, 0.09% NaCl). The saline was then collected and mounted on a slide and observed under an Olympus microscope equipped with X20 objective. The estrous cycle-stage was assessed according to epithelial cells morphology and the presence or absence of leukocytes. Mice that exhibited regular progress of their cycle for 3 consecutive days were eligible for further experiments.

### Ovaries extraction and handling

Mice were anesthetized for 2 minutes using isoflurane and were euthanized by cervical dislocation. Each mouse’s ovaries were extracted and washed in RPMI-1640 media (Sigma-Aldrich) containing 10% FBS and transferred to 1.5mL microcentrifuge tubes containing the same media. Next, the ovaries were cut using scissors and incubated for 30 minutes with ~7,800 IU/mL (Lot dependent) of collagenase type IV (Sigma-Aldrich) at 37°C. After which, 2.5μg/mL of DNase I (Sigma-Aldrich) was added, and tubes were incubated for additional 30 minutes. During incubation, the tubes were mixed every couple of minutes. Tissue homogenate was filtered through a 40μm strainer (Greiner Bio-One) and washed with 0.5mL RPMI media 5 times. Total cell count was then calculated using LUNA automated cell counter (Logos Biosystems). Cells were further processed for sorting (for single-cell RNA sequencing) or staining (for flow cytometry experiments).

### Cell sorting

Cell samples went through a series of centrifugations (400g, 5 minutes, at 4°C) and were stained with PE anti-CD45 (30-F11, BioLegend) in staining buffer (see flow cytometry section) for 30 minutes at 4°C. CD45-positive cells were sorted and collected using FACSAria™ III Cell Sorter (BD Biosciences). Collected samples were centrifuged and brought to a final volume of ~50μL, counted using LUNA automated cell counter (Logos Biosystems), and further processed for single-cell sample preparation.

### Single-cell RNA sequencing

Samples were prepared as outlined by the 10x Genomics Single Cell 3′ v2 Reagent Kit user guide. Briefly, the samples were washed twice in PBS (Sigma-Aldrich) + 0.04% BSA (Sigma-Aldrich) and re-suspended in PBS + 0.04% BSA. Sample viability was assessed with Trypan Blue (Biological Industries) using LUNA automated cell counter (Logos Biosystems) and validated using a hemocytometer. Following counting, the appropriate volume for each sample was calculated for a target capture of 10,000 and 7,700 cells (old and young samples, respectively) and loaded onto the 10x Genomics single-cell-A chip. After droplet generation, samples were transferred onto a 96-well plate, and reverse transcription was performed using a Veriti 96-well thermal cycler (Thermo Fisher). After the reverse transcription, cDNA was recovered using Recovery Agent provided by 10x Genomics followed by a Silane DynaBead clean-up (Thermo Fisher) as outlined in the user guide. Purified cDNA was amplified before being cleaned up using SPRIselect beads (Beckman Coulter). Samples were quantified with Qubit Fluorometer (Invitrogen) and run on Agilent TapeStation for quality control. cDNA libraries were prepared as outlined by the Single Cell 3′ Reagent Kits v2 user guide with appropriate modifications to the PCR cycles based on the calculated cDNA concentration (as recommended by 10X Genomics). Post library construction quantification and QC were performed as mentioned above for the post cDNA amplification step. Sequencing was performed with NextSeq500 system (Illumina), with ~50,000 reads per cell and 75 cycles for each read.

Seurat R package^64^ was used to read the 10X output data from Cell Ranger v3.0.1 according to the suggested protocol. Yield was 3,693 and 5,644 cells for young and old samples, respectively. Quality control tests were conducted to eliminate duplicates, and dead or low-quality cells – cells with less than 200 features, more than 2500 features, or more than 10% features of mitochondrial genes were excluded from further analysis. In total, we ended up with 3,307 and 5,468 cells for young and old samples, respectively (data available at Supplementary Tables 3 and 4).

### Cell type annotations

The Seurat R package, along with the SingleR package^65^ were used to analyze single-cell RNA sequencing data. After normalization of the data and clusters identification using Seurat’s graph-based clustering method, t-distributed stochastic neighbor embedding (tSNE) was used as a dimension reduction and visualization tool. Next, the SingleR algorithm was used to achieve the automatic annotation for each cell. Briefly, the algorithm compares each cell’s transcriptome to known transcriptomic “signatures” from reference genomes taken from The Immunological Genome Project (ImmGen)^66^. The algorithm calculates the correlation between the cell to different cell types, and based on the highest correlation suggests an annotation for the cell.

In addition, a set of literature-based gene markers was chosen to identify the different immune cell types. For each gene marker, a normalized score was calculated. *E*_*g,i*_ denotes the expression level of the gene *g* within the i^th^ cell; *M*_*g*_ and *m*_*g*_ denote the maximal and minimal expression of the gene *g* across all cells in the sample, respectively. *N*_c_ denotes the number of cells within cluster *c*, and *N*_*g,c*_ denotes the number of cells within cluster *c* that express the gene *g*. The normalized expression score, *S*_*g,i,c*_, for the gene *g* in the i^th^ cell within the cluster *c* is the multiplication of the gene relative expression and the relative fraction of cells within the cluster that express this gene, 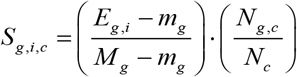. Figure 1c illustrates the distribution of the normalization score for a particular gene *g* over all the cells in cluster *c*. Normalized scores are between 0 to 1.

### Flow cytometry analysis

Ovarian cell suspensions were stained for flow cytometry analysis as the commercial protocol suggested (BD Biosciences). In short, cells were plated in 96-well U-shaped plates and went through a series of centrifugations (400g for 5 minutes, at 4°C) for media and debris cleaning. Next, samples were stained as the commercial protocol suggested using the following antibodies (purchased from BioLegend) diluted in staining buffer (PBS containing 0.09% sodium azide (Sigma-Aldrich)): PE anti-CD45 (30-F11), BV421 anti-CD11b (M1/70), APC/Cy7 anti-F4/80 (BM8), APC anti-CD3 (17A2), Pacific Blue anti-CD3 (145-2C11), APC/Cy7 anti-NK1.1 (PK136), PE/Cy7 anti-TCRβ (H57-597), Alexa Fluor 700 anti-CD8a (53-6.7) and FITC anti-CD4 (17A2). Staining was performed at 4°C for 30 minutes. Flow cytometry analyses were made using the S100EXi (Stratedigm) flow cytometer. Viability test of CD45^+^ cells was conducted using Zombie-NIR (BioLegend).

### Statistical analyses

All statistical analyses were calculated using MATLAB R2019b (MathWorks). We used 2-samples Kolmogorov-Smirnov test (two-tailed, p-value <0.05) for identifying significant changes in CD3^+^ lymphocytes and macrophages fractions at different ages. We set a threshold of 2-fold change and used student’s t-test (two-tailed, p-value <0.05) For defining DEGs. We used χ^2^ test for frequencies comparison (two-tailed, p-value <0.05) while setting a threshold of 10% difference for significant fraction differences in gene expression.

### GO enrichment analysis

Significant downregulated or upregulated genes for each cell type were taken for GO enrichment analysis of biological processes using the g:Profiler platform^67^ (database versions: Ensembl 104, Ensembl Genomes 51, and Wormbase ParaSite 15, published on 5/26/2021). Next, all significant processes were further analyzed using the REVIGO platform for eliminating redundant GO terms^68^. Then, cell-type specific and global (for at least two cell types) processes were found and analyzed once more using the REVIGO platform in order to cluster them according to their semantics distance.

### Cytokines and chemokines networks

The Kyoto Encyclopedia of Genes and Genomes (KEGG) was used to construct the chemokines and cytokines ligand-receptor networks. These were built based on the Cytokine-cytokine receptor interaction - Mus musculus (mouse) map (pathway map mmu04060)^69^.

## Supporting information

Supplementary infromation

## Data availability

Source data are provided with this paper

