## Supplementary infromation for "Aged mouse ovarian immune milieu shows a shift towards adaptive immunity and attenuated cell function"

### Supplementary Information

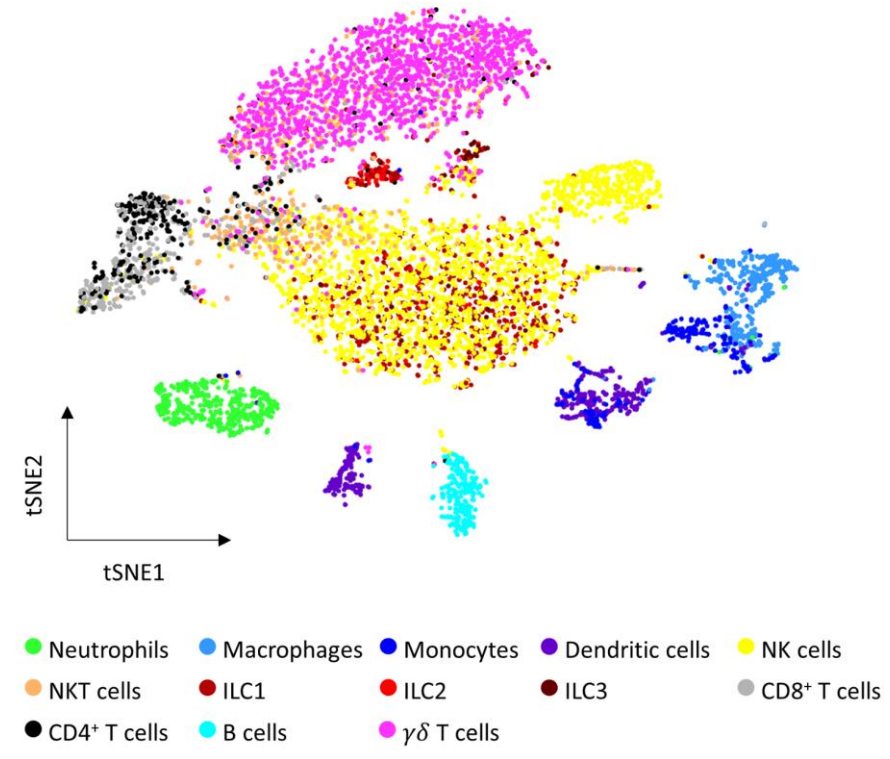


**Supplementary Figure 1: Ovarian immune cells annotation.** tSNE plot of both young and old samples annotated using the SignleR algorithm[1].

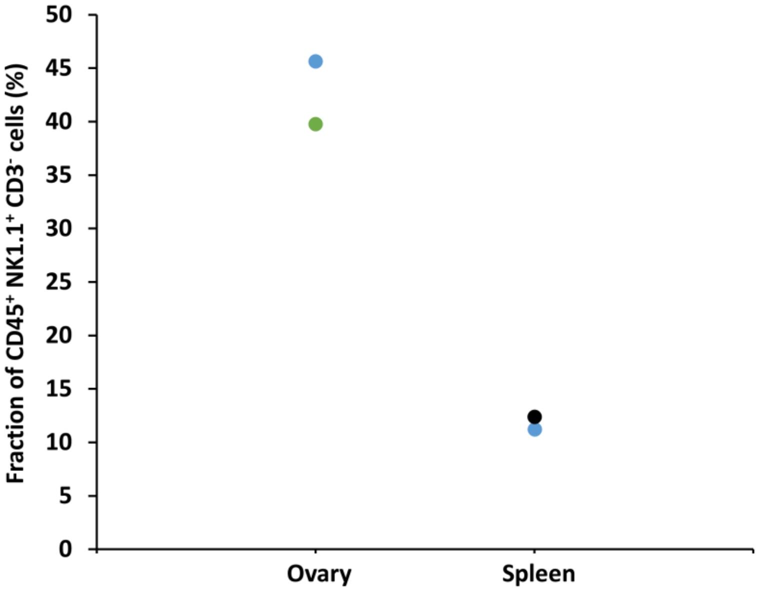


**Supplementary Figure 2: Flow cytometry measurement of group 1 innate lymphoid cells (G1-ILCs) fraction.** Young female mice's (*n*=3, 12 weeks old) immune cells were collected from the ovaries and spleen and were analyzed using flow cytometry for their group 1 innate lymphoid cells ($CD45^+$ $NK1.1^+$ $CD3^-$) distribution. Each color represent a different mouse.

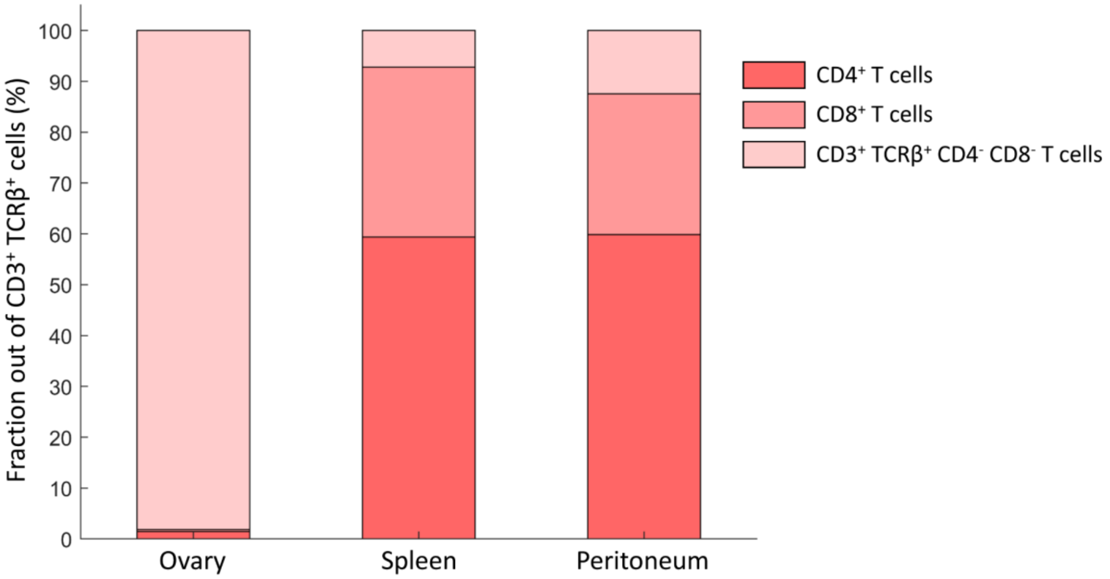


**Supplementary Figure 3: Double-negative T cells are more abundant in the ovaries compared to other tissues.** Old female mouse's (55 weeks) immune cells were collect from the ovaries, spleen and peritoneum and were analyzed using flow cytometry for their T-lymphocytes distribution. While most cells in spleen and peritoneum were $CD4^+$ and $CD8^+$ T cells as anticipated, almost all T cells in the ovaries were $CD4^-$ $CD8^-$.

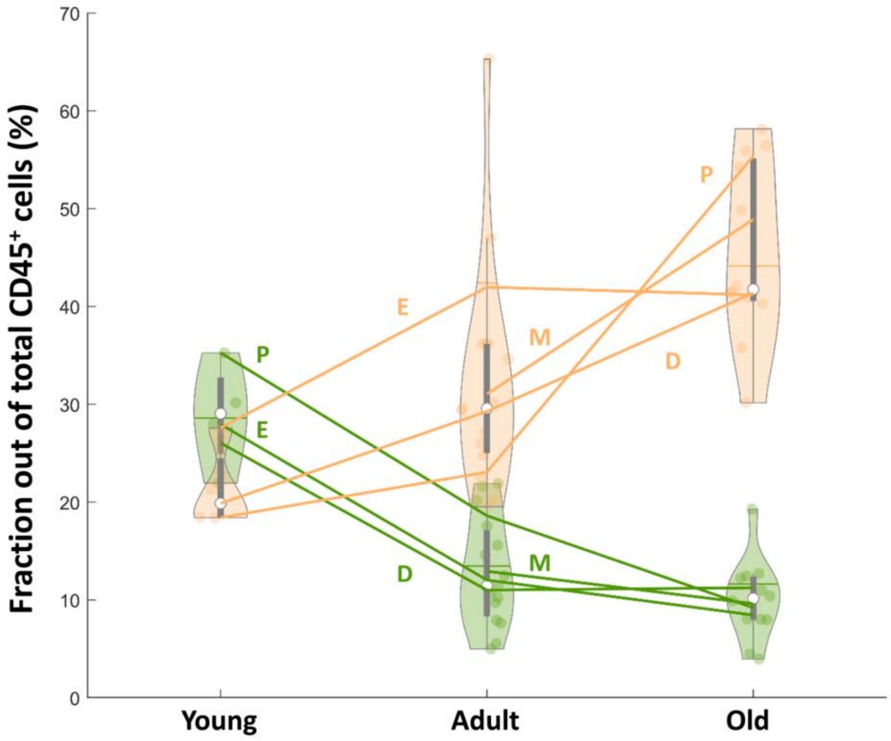


**Supplementary Figure 4: Changes in ovarian CD3+ cells and macrophages is cycle stage independent.** Violin plot of macrophages and CD3+ lymphocytes fractions out of total CD45+ as shown in Fig. 2D with an additional annotation of the cycle stage. Lines represent the mean fraction of each population in each stage of the estrus cycle (P – Proestrus; E – Estrus; M – Metestrus; D - Diestrus). $N_{Young\_P} = 1$, $N_{Young\_E} = 1$, $N_{Young\_D} = 2$, $N_{Adult\_P} = 2$, $N_{Adult\_E} = 4$, $N_{Adult\_m} = 4$, $N_{Adult\_D} = 5$, $N_{Old\_P} = 2$, $N_{Old\_E} = 2$, $N_{Old\_M} = 3$, $N_{Old\_D} = 5$;

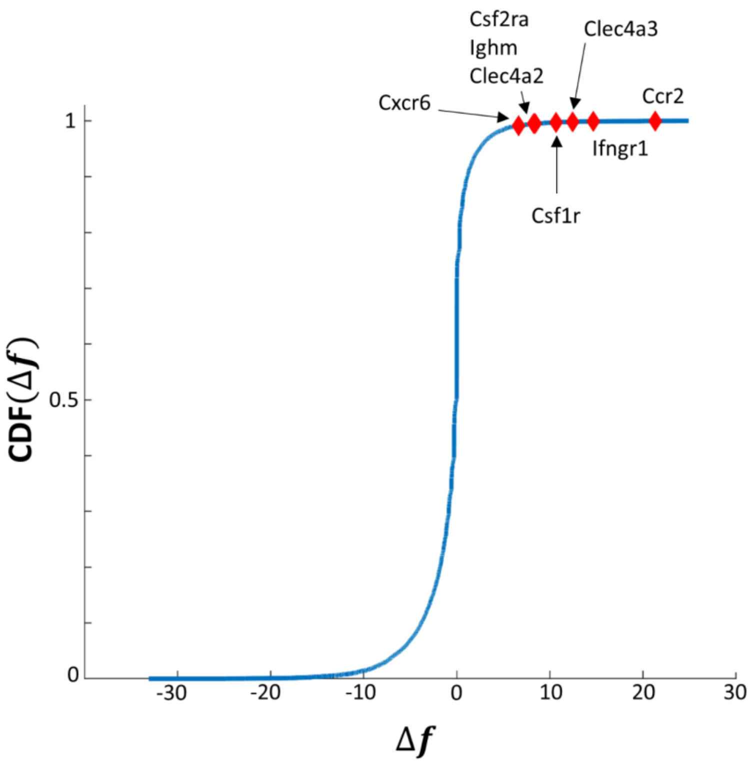


**Supplementary Figure 5: Increase in recognition of senescent cells-related genes in old ovarian macrophages.** $\Delta f = f_{old} - f_{young}$ is the difference between the fraction of old and young macrophages that expresses the relevant genes. The cumulative distribution function (CDF) of changes in the expression fraction of each gene at old age. The probability of getting at random $\Delta f$ that is higher than the senescent cell recognition genes' $\Delta f$ is less than 0.01.
